# Le Petit Prince Hong Kong (LPPHK): Naturalistic fMRI and EEG data from older Cantonese speakers

**DOI:** 10.1101/2024.04.24.590842

**Authors:** Mohammad Momenian, Zhengwu Ma, Shuyi Wu, Chengcheng Wang, Jonathan Brennan, John Hale, Lars Meyer, Jixing Li

## Abstract

Currently, the field of neurobiology of language is based on data from only a few Indo-European languages. The majority of this data comes from younger adults neglecting other age groups. Here we present a multimodal database which consists of task-based and resting state fMRI, structural MRI, and EEG data while participants over 65 years old listened to sections of the story The Little Prince in Cantonese. We also provide data on participants’ language history, lifetime experiences, linguistic and cognitive skills. Audio and text annotations, including time-aligned speech segmentation and prosodic information, as well as word-by-word predictors such as frequency and part-of-speech tagging derived from natural language processing (NLP) tools are included in this database. Both MRI and EEG data diagnostics revealed that the data has good quality. This multimodal database could advance our understanding of spatiotemporal dynamics of language comprehension in the older population and help us study the effects of healthy aging on the relationship between brain and behaviour.

## Background & Summary

Language comprehension in daily life is a process which does not happen in a vacuum; the environment could be noisy, people have different speaking rates (some speak faster than others), and sociocultural variables such as social class might influence what variety of language people speak. Most of the research in the neurobiology of language is done on linguistic processes which are not representative of the language used in natural contexts. There is, however, an increasing interest in doing research using more naturalistic tasks (See Supplementary Table 1 for a list of published databases on naturalistic language). Naturalistic data could be more reliable and ecologically valid because such data is representative of what the brain does in real life^1^. Using naturalistic data, researchers can study language at different levels of language (word, phrase, sentence, and discourse) at the same time and investigate how these levels interact with each other^2^. Naturalistic tasks allow immense flexibility in the study of special populations such as, for instance, people with autism spectrum disorder (ASD) where sticking to the experimental protocol could be really challenging.

Language comprehension is usually studied using techniques such as fMRI and M/EEG. fMRI has an advantage in spatial resolution showing where exactly in the brain a mental process happens. However, in studying language temporal dynamics are as important as spatial correlates. MEG and EEG have great temporal resolution making them valuable tools for studies of language comprehension. Almost all previous naturalistic databases have used fMRI neglecting to cater for questions about temporal dynamics of language^3^ (see Supplementary Table 1). There is a need for naturalistic studies which provide data on spatiotemporal dynamics of language.

Research shows that the majority of previous studies in psycholinguistics (85%) are done in only 10 languages with English accounting for 30% of the published studies^4^. Most Indo-European languages are very similar in terms of morpho-syntactic properties and script. More research needs to be done on understudied languages which have different properties. This could accommodate more diversity in the research on neurobiology of language and help the findings be more representative of the world’s population.

One issue with prior naturalistic research is that they have only provided data on healthy younger adults (See Supplementary Table 1) neglecting other populations such as children and older adults. The World Population Aging Report^5^ shows that the population of seniors aged 65 years and over will double in the next three decades. This means that all stakeholders need to be ready to embrace this population shift. It will be, therefore, very important to study how healthy aging could influence the relation between brain and behavior, particularly in naturalistic settings where the cognitive demands could be higher for older people.

We know that healthy aging is usually accompanied by individual differences in cognitive decline^6^. Any database on language in older adults should also provide information about key cognitive skills which are involved in language comprehension if resources are available. Without data on cognitive functions, it will be difficult to document the role of individual differences in language comprehension in healthy aging. Only one database (see^1^) has provided such data.

In this study, we present a database which includes 52 Cantonese speakers (over 65 years old) listening to 20 minutes of the story The Little Prince. We collected both fMRI (task based and resting state) and EEG data from the same participants. Data on language history and lifetime experiences as well as behavioural data about linguistic skills and cognitive functions such as inhibition, short-term memory, processing speed, verbal fluency, and picture naming speed were collected. We think this is the first naturalistic database on older people which provides both fMRI and EEG data as well as extensive cognitive data for each participant. This valuable contribution aligns with our vision for the LPP database expansion^7^, incorporating diverse modalities, languages, and annotations to foster interdisciplinary research in psycho- and neurolinguistics.

## Methods

### Participants

We recruited 52 healthy, right-handed older Cantonese participants (40 females, mean age=69.12, SD=3.52) from Hong Kong for the experiment, which consists of an fMRI and an EEG session. In both sessions, participants listened to the same sections of *The Little Prince in Cantonese* for approximately 20 minutes. We made sure each participant was right-handed and a native Cantonese speaker using the Language History Questionnaire^8^ (LHQ3). Additionally, participants reported normal or corrected normal hearing. They confirmed they had no cognitive decline. Two participants did not take part in the fMRI session and an additional 4 participants’ fMRI data were removed due to excessive head movement, resulting in a total of 46 participants (39 females, mean age=69.08yrs, SD=3.58) for the fMRI session and 52 participants (40 females, mean age=69.12yrs, SD=3.52) for the EEG session. Supplementary Table 2 shows participants’ demographic information and quiz response accuracy. Prior to the experiment, all participants were provided with written informed consent. All participants received monetary compensation after each session. Ethical approval was obtained from the Human Subjects Ethics Application Committee at the Hong Kong Polytechnic University (application number HSEARS20210302001). This study was performed in accordance with the Declaration of Helsinki and all other regulations set by the Ethics Committee.

### Procedures

The study consisted of an fMRI session and an EEG session. The order of the EEG and fMRI sessions was counterbalanced across all participants, and a minimum two-week interval was maintained between sessions.

#### fMRI experiment

Before the scanning day, an MRI safety screening form was sent to the participants to make sure MRI scanning was safe for them. We also sent them simple readings and videos about MRI scanning so that they could have an idea of what it would be like to be in a scanner. On the day of scanning, participants were initially introduced to the MRI facility and comfortably positioned inside the scanner, with their heads securely supported using paddings. An MRI-safe headphone was provided for participants to wear inside the head coil. The audio volume for the listening task was adjusted to ensure audibility for each participant. A mirror attached to the head coil allowed participants to view the stimuli presented on a screen. Participants were instructed to stay focused on the visual fixation sign while listening to the audiobook.

The scanning session commenced with the acquisition of structural (T1-weighted) scans. Subsequently, participants engaged in the listening task concurrently with fMRI scanning. The task-based fMRI experiment was divided into four runs, each corresponding to a section of the audiobook. Comprehension was assessed by a series of 5 yes/no questions (20 questions in total, see Supplementary Table 2) on the content they had listened to. These questions were presented on the screen, with participants indicating their answers by pressing a button. The session concluded with the collection of resting-state fMRI data.

#### Cognitive tasks

Four cognitive tasks were selected to assess participants’ cognitive abilities in various domains, including the digit span forward task, picture naming task, verbal fluency task, and Flanker task (see Figure 2). These tasks were delivered after the fMRI session in a separate soundproof booth.

**Fig. 1.**
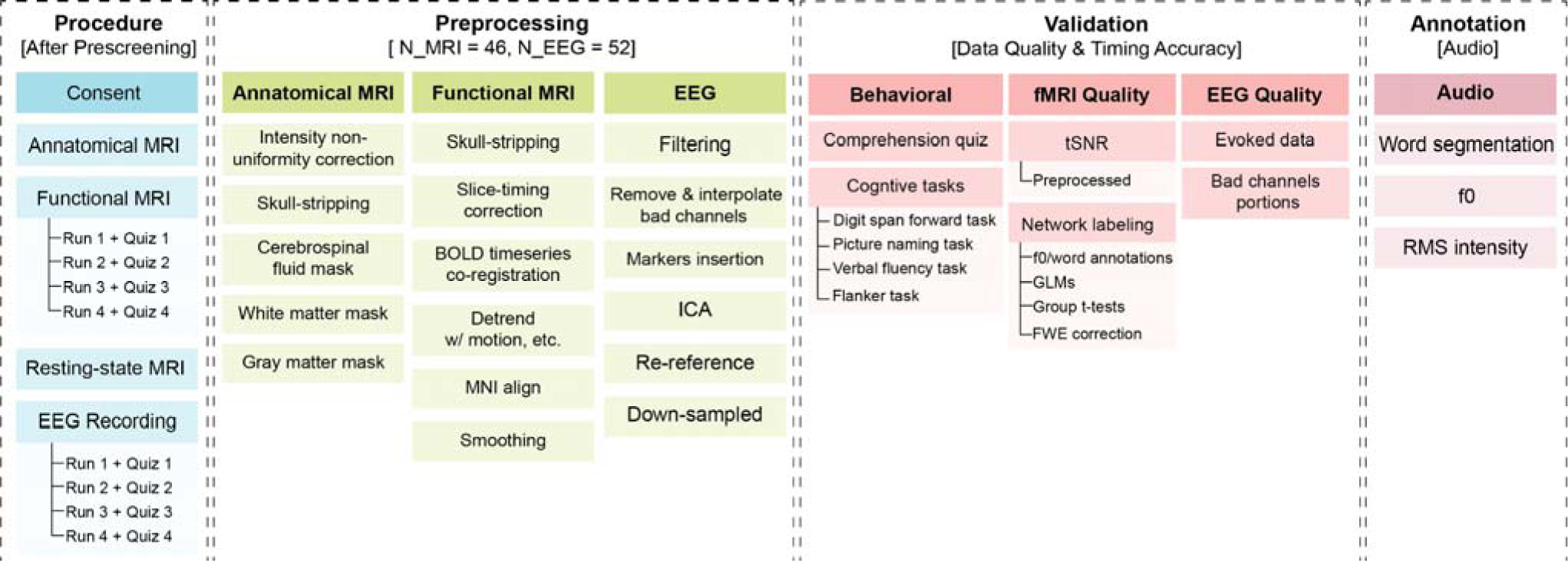
Schematic overview of the LPPHK data collection procedures, preprocessing, technical validation and annotation. During data collection (blue), neural imaging signals were measured while participants listened to 4 sections of the audiobook. The sessions were counterbalanced with half of the participants starting with fMRI and the other half with EEG, The sequence for fMRI data collection always commenced with anatomical MRI, followed by functional and resting-state MRI. After preprocessing the data (green), behavioral and overall data quality were assessed (light red). Audio and text annotations were extracted using NLP tools (dark red)

**Fig. 2.**
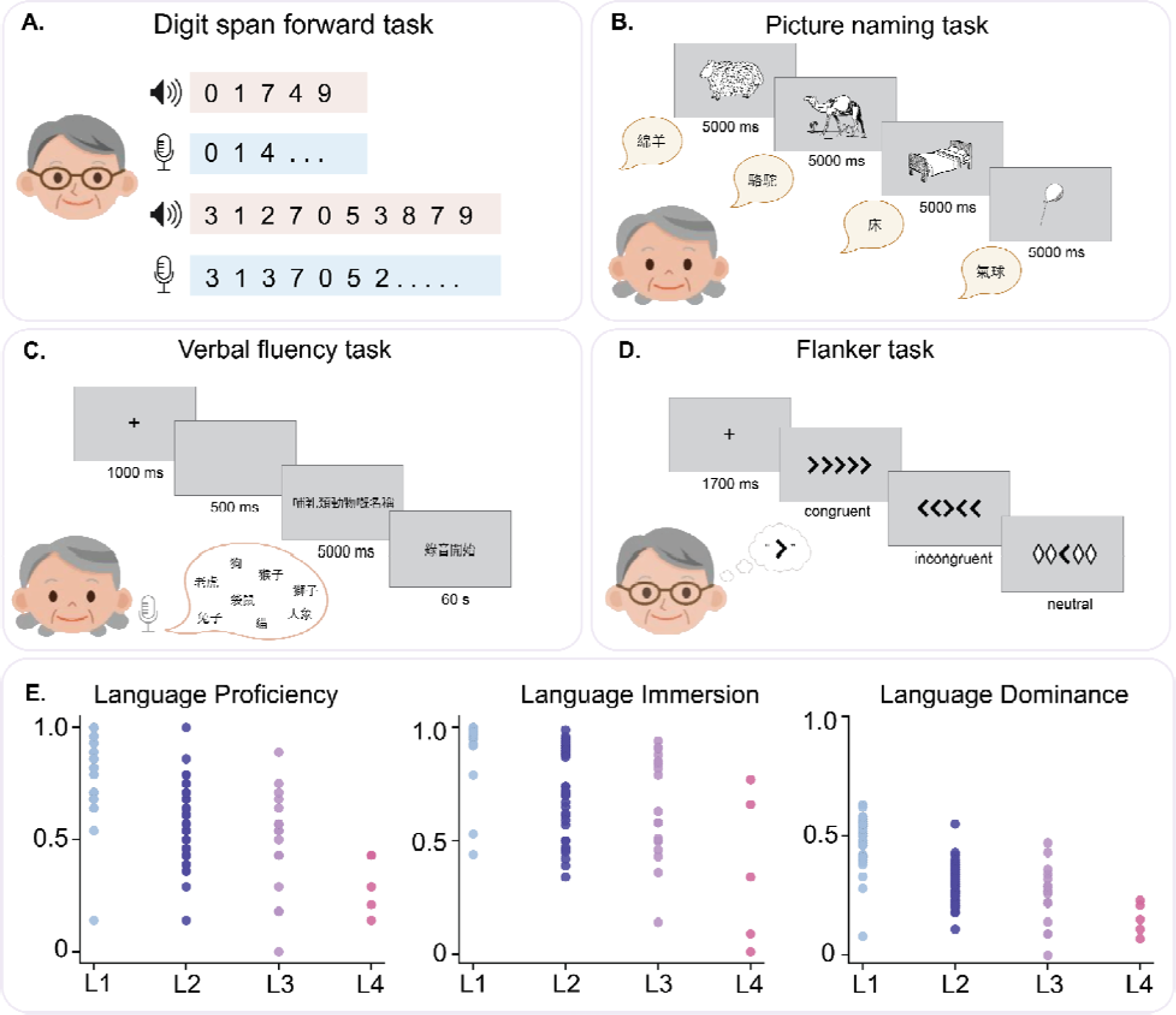
Behavioral cognitive tasks procedure and Language History Questionnaire (LHQ) score distribution. **(A)** Digital span forward task. **(B)** Picture naming task. **(C)** Verbal fluency task. **(D)** Flanker task. **(E)** Language History Questionnaire (LHQ) provides scores for proficiency, immersion, and dominance for the languages participants reported. All the participants can speak Cantonese, and only five can speak four languages.

The digit span forward task^9^ measured the span of short-term memory. Participants were presented with audible random sequences of digits 0-9 in Cantonese and were instructed to verbally recall them in the same order (see Fig. 2A). The participants had a chance to practice before the main task started. The span ranged from 4 digits to 14 digits. For scoring, we gave partial credit (0.5) if participants could answer 1 item correctly and full credit (1) for 2-3 correct answers (See Supplementary Table 3).

In the picture-naming task, participants were required to quickly and accurately name objects and actions using line drawings from the Object and Action Naming Battery^10^ (ONAB). This battery has been normed across multiple languages, including Cantonese^11^ (see Fig. 2B). Participants had a brief practice session before the main task started. Average picture naming latency is reported in Supplementary Table 3 for each participant.

The verbal fluency task^9^ involved participants generating as many words as possible within 60 seconds that fit into four randomized test categories (mammals, non-mammals, fruits, electronic appliances). Verbal responses were recorded and later scored by independent judges (see Fig. 2C). The score is the average score among all four categories (See Supplementary Table 3).

The Flanker task^12^ assessed response inhibition and processing speed. Participants were asked to identify the direction of a central target arrow while disregarding four flanking arrows (see Fig. 2D). The score for inhibition is usually calculated as a difference score where the latency in the congruent condition is subtracted from the latency in the incongruent condition. We presented raw RT for each condition in Supplementary Table 3 so that researchers could use them flexibly based on their research questions. We considered participants’ average reaction time in neutral conditions as an indication of processing speed (see Supplementary Table 3). In the neutral condition, there are no conflicting or facilitating cues which makes it a good measurement for processing speed^13^.

#### EEG experiment

During the EEG experiment, participants were seated comfortably in a quiet room and standard procedures were followed for electrode placement and EEG cap preparation. Participants were instructed to focus on a fixation sign displayed on a monitor. The EEG recording was then initiated, with participants listening to the audiobook. The audio volume was adapted to each participant’s hearing ability before the recording using a different set of stimuli. Similar to the fMRI experiment, participants listened to four sections of the audiobook, each lasting approximately 5 minutes. After each run, participants were asked to answer a total of 20 yes/no questions, with 5 questions assigned to each run. They indicated their answers by pressing a button. The EEG recording was conducted continuously throughout all four runs until their completion.

#### Questionnaires

We administered LHQ3 and the Lifetime of Experiences Questionnaire^14^ (LEQ) during EEG cap preparation. The participants did not need to move or fill in these questionnaires themselves; a research assistant asked the questions one by one in Cantonese and input the responses in an online Google form. LHQ is designed to document language history by producing aggregate scores for language proficiency, exposure, and dominance in all the languages spoken by the participants (See Fig. 2E). LEQ is a tool to document what sorts of activities (e.g. sports, music, education, profession, etc) participants engage in over their lifetime. It measures lifetime experiences in three periods of life: from 13 to 30 (young adulthood), from 30 to 65 (midlife), and after 65 (late life). LEQ produces a total score (see participants.tsv in the OpenNeuro) which is an indication of cognitive activity. Collecting data using these two questionnaires allowed us to have a thicker description of our participants’ linguistic, social, and cognitive experiences.

### Stimuli

The experimental stimuli utilized in both the EEG and fMRI consisted of approximately 20 minutes of the story *The Little Prince in Cantonese* audiobook. It was translated and narrated in Cantonese by a native male speaker. The stimuli consist of a total of 4,473 words and 535 sentences. To facilitate data analysis and participant engagement, the stimuli were further segmented into four distinct sections, each spanning nearly five minutes. To assess listening comprehension, participants were presented with five yes/no questions after completing each section, resulting in a total of 20 questions throughout the experiment. To make sure the speed of story narration was normal for the participants, we asked a few people who were different from the participants in this study to judge the speed and comprehensibility. They all reported the speed was normal, neither so slow nor so fast.

### Data acquisition

The MRI data were collected at the University Research Facility in Behavioral and Systems Neuroscience (UBSN) at The Hong Kong Polytechnic University. EEG data was collected at the Speech and Language Sciences Laboratory within the Department of Chinese and Bilingual Studies at the same university.

#### fMRI data

MRI imaging data were acquired using a 3T Siemens MAGNETOM Prisma system MRI scanner with a 20-channel coil. Structural MRI was acquired for each participant using a T1-weighted sequence with the following parameters: repetition time (TR) = 2,500 ms, echo time (TE) = 2.22 ms, inversion time (TI) = 1,120 ms, flip angle α (FA) = 8°, field of view (FOV) = 240 × 256 × 167 mm, resolution = 0.8 mm isotropic, acquisition time = 4 min and 32s. The acquisition parameters for echo planar T2-weighted imaging (EPI) were as follows: 60 oblique axial slices, TR = 2000 ms, TE = 22 ms, FA= 80°, FOV = 204 × 204 × 165 mm, 2.5 mm isotropic, and acceleration factor 3. E-Prime 2.0 (Psychology Software Tools) was used to present the stimuli.

#### EEG data

A 64-channel Neuroscan system on a 10-20 electrode template was used for data acquisition, sampling at a rate of 1000 Hz. To mark the onset of each sentence, triggers were set at the beginning of each sentence. STIM2 software (Compumedics Neuroscan) was used for stimulus presentation.

### Preprocessing

All MRI data were preprocessed using the *NeuroScholar* cloud platform (http://www.humanbrain.cn, Beijing Intelligent Brain Cloud, Inc.), provided by The Hong Kong Polytechnic University. This platform uses an enhanced pipeline based on *fMRIPrep^15,16^ 20.2.6* (RRID: SCR_016216) and supported by *Nipype^17^ 1.7.0* (RRID: SCR_002502).

#### Anatomical

The structural MRI data underwent intensity non-uniformity correction^18^, skull-stripping, and brain tissue segmentation of cerebrospinal fluid (CSF), white matter (WM), and gray matter (GM) based on the reference T1w image. The resulting anatomical images were nonlinearly aligned to the ICBM 152 Nonlinear Asymmetrical template version 2009c (MNI152NLin2009cAsym) template brain (see Fig. 1).

#### Functional

The preprocessing of both resting and functional MRI data included the following steps: (1) skull-stripping, (2) slice-timing correction with the temporal realignment of slices according to the reference slice, (3) BOLD time-series co-registration to the T1w reference image, (4) head-motion estimations and spatial realignment to adjust for linear head motion, (5) applying parameters from structural images to spatially normalize functional images into Montreal Neurological Institute (MNI) template, and (6) smoothing by a 6mm FWHM (full-width half-maximum) Gaussian kernel.

#### EEG

The pre-processing was carried out using EEGLAB^19^ and in-house MATLAB functions. The preprocessing of EEG data included the following steps: (1) a cutoff frequency filter with 1 Hz high pass and 40.0 Hz low pass cut-off was applied followed by a notch filter at 50 Hz to reduce electrical line noise, (2) use of kurtosis measure to identify and remove bad channels, (3) application of the RUNICA algorithm (from EEGLab toolbox, 2023 version), a machine learning algorithm that evaluates ICA-derived components, for automated rejection of artifacts, including signal noise from eye and muscle, high-amplitude artifacts (e.g., blinks), and signal discontinuities (e.g., electrodes losing contact with the scalp), (4) interpolating data for bad channels using spherical splines for each segment. (5) re-referencing the data by using both electrodes M1 and M2 as the reference for all channels and (6) down-sampling all the data to 250 Hz.

### Annotations

We present audio and text annotations, including time-aligned speech segmentation and prosodic information, as well as word-by-word predictors derived from natural language processing (NLP) tools. These predictors include aspects of lexical semantic information, such as part-of-speech (POS) tagging and word frequency. An overview of our annotations is presented in fig. 3. These annotations can also be accessed on OpenNeuro (see the Data Records section for details).

**Fig. 3.**
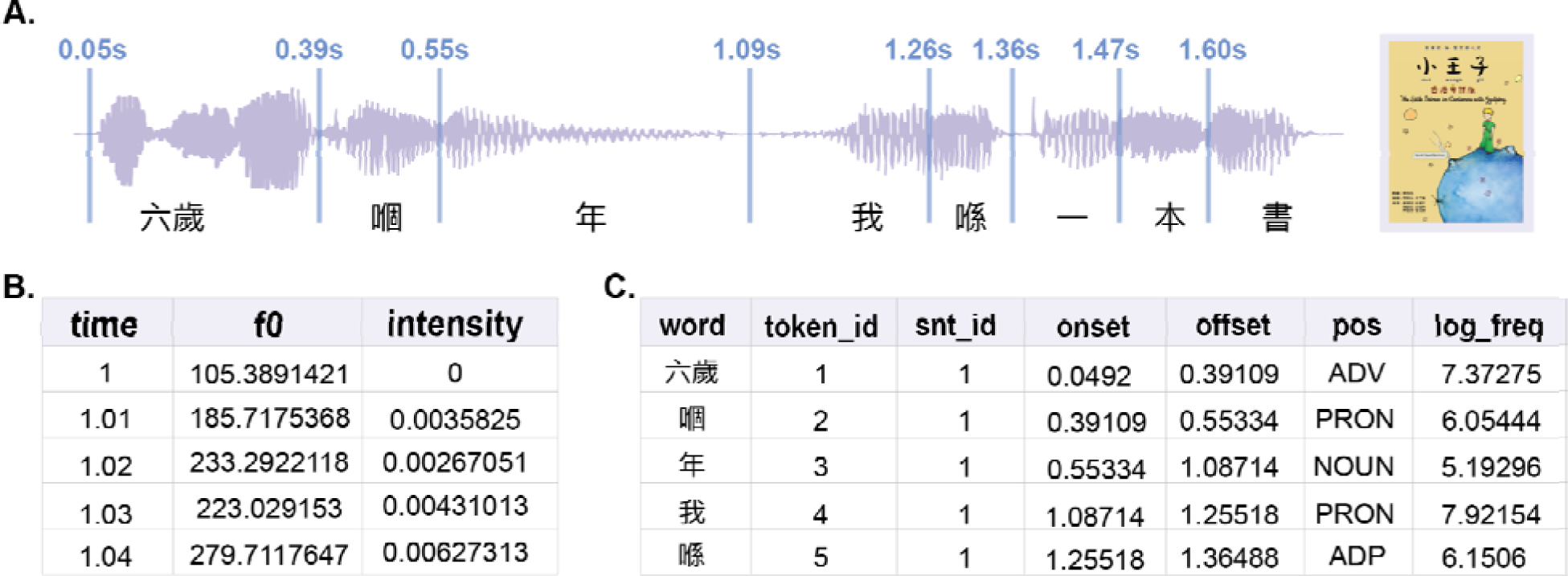
Annotation information for the stimuli. (**A**) Word boundaries in the audio files, included in files: lppHK_word_information.txt. (**B**) f0 and RMS intensity for every 10 ms of the audios, included in files: wav_acoustic.csv (**C)** Tokenization, POS tagging, and log-transformed word frequency in files: lppHK_word_information.txt.

#### Prosodic information

We extracted the root mean square intensity and the fundamental frequency (f0) from every 10 ms interval of the audio segments by utilizing the Voicebox toolbox (http://www.ee.ic.ac.uk/hp/staff/dmb/voicebox/voicebox.html). Peak RMS intensity and peak f0 for each word in the naturalistic stimuli were used to represent the intensity and pitch information for each word.

#### Word frequency

Word segmentation was performed manually by two native Cantonese speakers. The log-transformed frequency of each word was also estimated using PyCantonese^20^, Version 3.4.0 (https://pycantonese.org/). The built-in corpus in PyCantonese is the Hong Kong Cantonese Corpus^21^ (HKCancor), collected from transcribed conversations between March 1997 and August 1998.

#### Part-of-speech tagging

Part-of-speech (POS) tagging for each word in the stimuli was extracted using the PyCantonese^20^, Version 3.4.0 (https://pycantonese.org/). Following the manual segmentation of words, we input these segments into the Cantonese-exclusive NLP tool PyCantonese, which then provided POS tags for each word according to the Universal Dependencies v2 tagset^22^ (UDv2).

### Data Records

We have anonymized all records by removing information and anatomical data that could potentially identify the participants. Results files are available from the OpenNeuro repository (https://openneuro.org/datasets/ds004718). For a visual representation of the data collection structure, refer to Fig. 4. A comprehensive description of the available content is provided in the README file on the repository. The scripts utilized for this research paper are accessible both on the repository and GitHub at https://github.com/jixing-li/lpp_data.

**Fig. 4.**
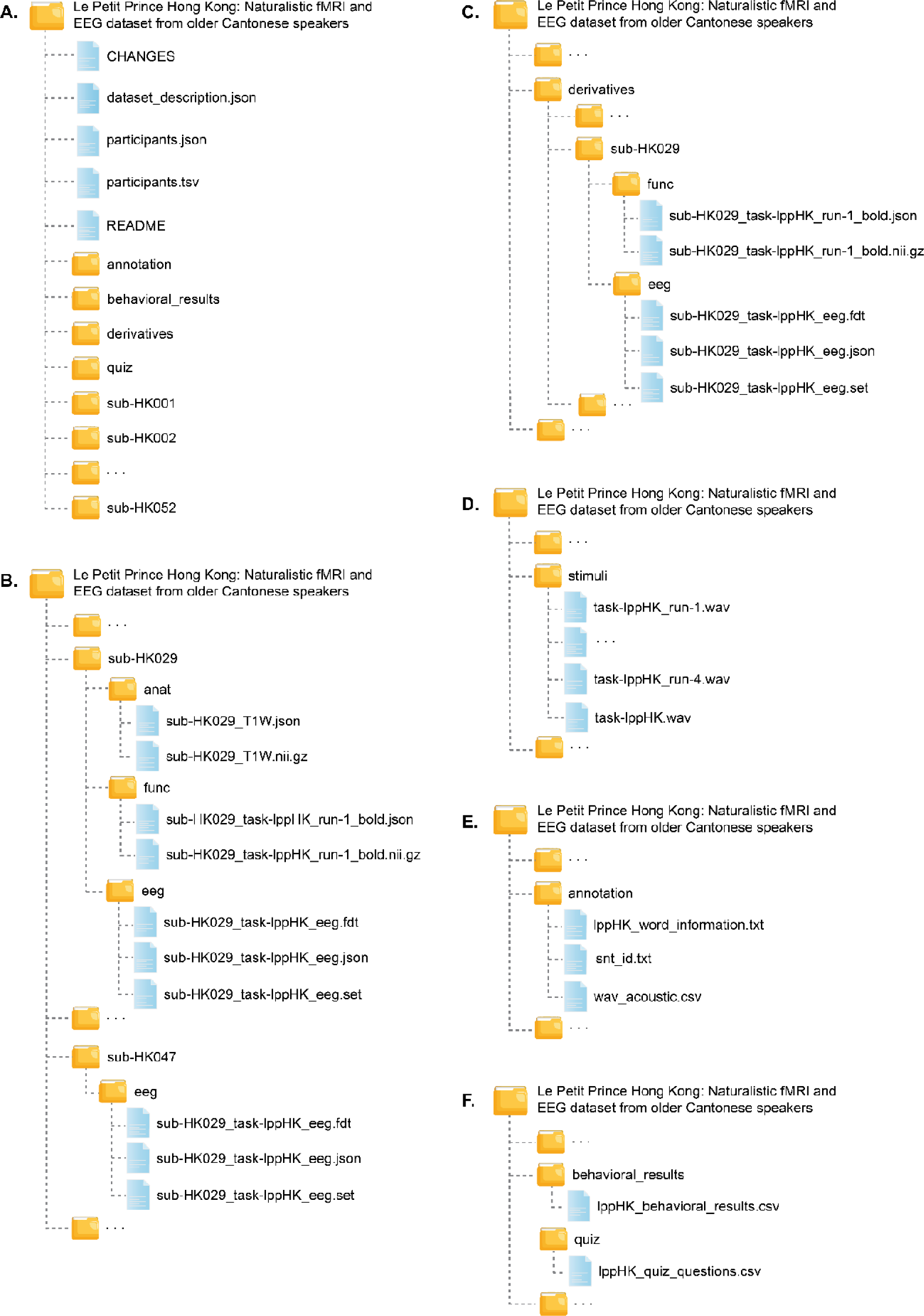
Organization of the data collection. **(A)** General overview of directory structure. **(B)** Content of subject-specific directories for raw data, including EEG, structural, resting-state MRI and functional MRI data. **(C)** Content of subject-specific directories of preprocessed data for both EEG and fMRI modals. **(D)** Content of the experimental stimuli directory. **(E)** Content of the annotation file directory. **(F)** Content of the quiz and cognitive behavioral task results directories.

**Participant responses.** Location participants.json, participants.tsv.

**File format** tab-separated value.

Participants’ sex, age, and accuracy of quiz questions for each fMRI and EEG experiment in tab-separated value (tsv) files. Data is structured as one line per participant.

**Audio files.** Location stimuli/task-lppHK_run-1[2-4].wav.

**File format** wav.

The 4-section audiobook from *The Little Prince* in Cantonese.

**Anatomical data files.** Location sub-HK<ID>/anat/sub-HK<ID>_T1w.nii.gz

**File format** NIfTI, gzip-compressed.

The raw high-resolution anatomical image after defacing.

**Functional data files.**

Location sub-HK<ID>/func/sub-HK<ID>_task-lppHK_run-1[2–4]_bold .nii.gz

**File format** NIfTI, gzip-compressed.

**Sequence protocol**

sub-HK<ID>/func/sub-HK<ID>_task-lppHK_run-1[2–4]_bold.json.

The preprocessed data are also available as derivatives/sub-HK<ID>/func/sub-HK<ID>_task-lppHK_run-1[2–4]_bold.nii.gz.

**Resting-state MRI data files.**

Location sub-HK<ID>/func/sub-HK<ID>_task-rest_bold.nii.gz

**File format** NIfTI, gzip-compressed.

**Sequence protocol**

sub-HK<ID>/func/sub-HK<ID>_task-rest_bold.json.

The preprocessed data are also available as derivatives/sub-HK<ID>/func/sub-HK<ID>_rest_bold.nii.gz.

**EEG data files.**

Location sub-HK<ID>/eeg/sub-HK<ID>_task-lppHK_eeg.set

**File format** set (a type of MATLAB file, with a file in the .fdt extension containing raw data)

The preprocessed data are also available as derivatives/sub-HK<ID>/eeg/sub-HK<ID>_task-lppHK_eeg.set(together with a file in the .fdt extension containing raw data)

**Annotations.** Location annotation/snts.txt, annotation/lppHK_word_information.txt, annotation/wav_acoustic.csv

**File format** comma-separated value.

Annotation of speech and linguistic features for the audio and text of the stimuli.

**Quiz questions.** Location quiz/lppHK_quiz_questions.csv

**File format** comma-separated value.

The 20 yes/no quiz questions were employed in both the fMRI and EEG experiments.

## Technical Validation

To ensure adequate comprehension, we calculated participants’ response accuracy of the quiz questions during the EEG and fMRI experiments. Quality assessment for EEG data was focused on the bad channel portions and the highest amplitude of epoched data. The quality of fMRI scans was evaluated through the computation of temporal signal-to-noise ratio (tSNR). Moreover, three whole-brain functional analyses were conducted utilizing pitch (f0), intensity and word rate annotations. These analyses serve to showcase the high data quality akin to prior research, while substantiating the precise timing accuracy between fMRI time-series for each participant^23,24^.

### Behavioral results

In both the EEG and fMRI experiments, participants were presented with the same set of 20 yes/no quiz questions in total. The average accuracy across participants was 66.25% (SD = 0.13) for the EEG experiment and 59.24% (SD = 0.12) for the fMRI experiment. The observed lower accuracy compared to a previous database^23^ could potentially be attributed to the advanced age range of our participants, as age-related memory decline may influence their performance. One sample question is shown below.

Example

Question: Did the Little Prince come to Earth from another planet?

Answer: Yes

### Quality assessment for EEG data

The quality assessment was conducted after preprocessing of EEG data. An overview of both qualitative and quantitative measurements for the signal quality is shown in Figure 5.

**Fig. 5.**
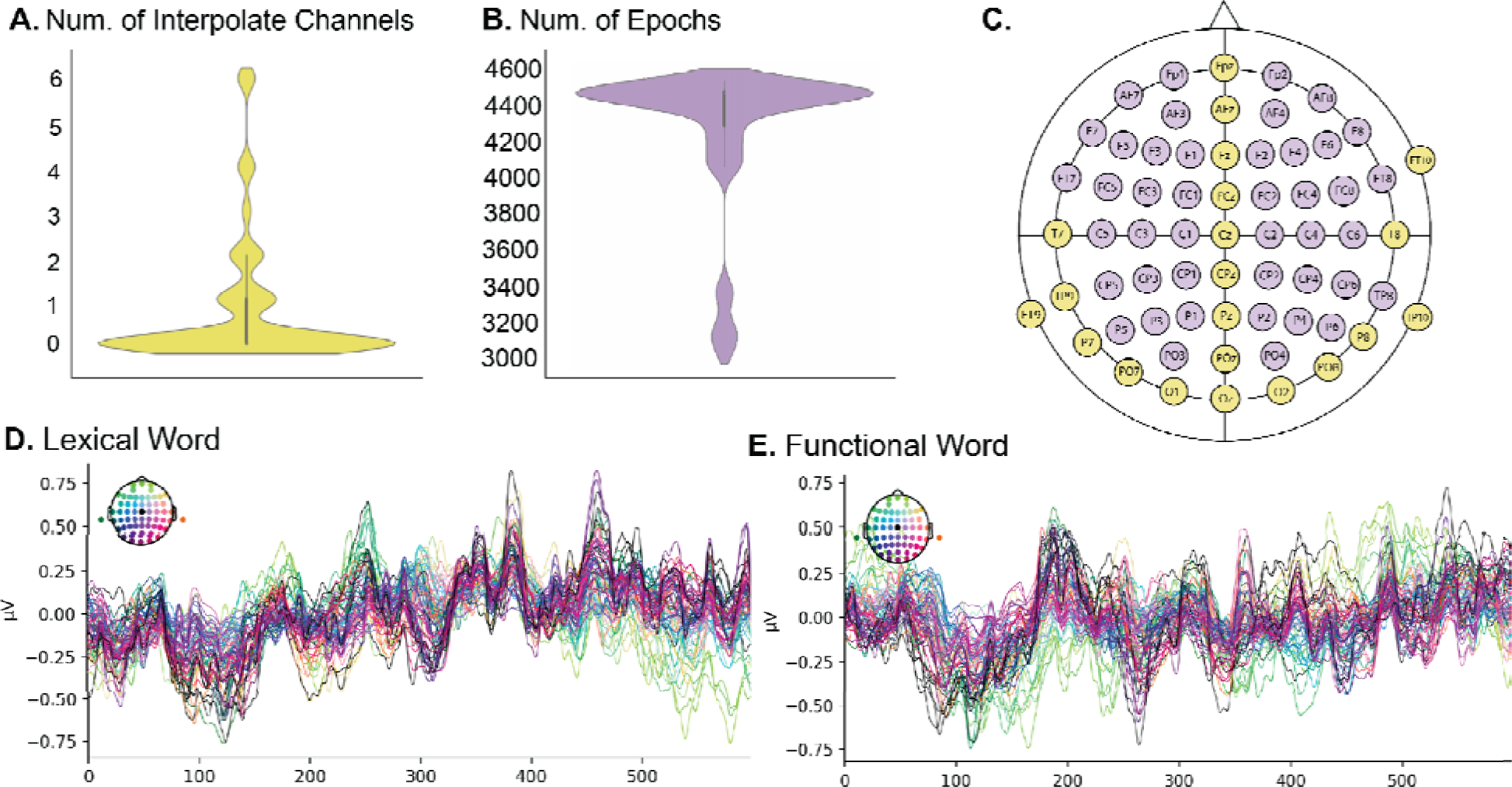
EEG Data quality assessment. **(A)** Number of interpolated channels per participant. **(B)** Number of epochs that survived from thresholded for each participant. **(C)** Sensor layout of the EEG system (64 channels, Neuroscan). The average response across 50 trials for both lexical words **(D)** and functional words **(E)**.

#### Number of Bad channel removals

By using the kurtosis measure, we identified bad channels and computed the number of interpolated channels, which is presented in Supplementary Table 4 and Fig. 5A. It is shown that only 15 out of 50 participants had bad channels during data acquisition which accounts for lower than 10% of all the channels.

#### High amplitudes of Epochs

Epochs with amplitudes of above +150 mV and below -150 mV in any of the 60 channels were excluded from averaging. Under the threshold of 150 mv, more than half of event epochs were rejected for four participants (incl. sub-HK005, sub-HK019, sub- HK029 and sub-HK037). We adjusted different thresholds for these participants for further analyses, with 300mV for sub-HK005, 200mV for both sub-HK019 and sub-HK037, and 350mV for sub-HK029. The epochs that survived from the threshold are counted and shown in Supplementary Table 4 and Fig. 5B.

### Event-related potentials (ERPs)

ERPs refer to brain activities that are elicited in response to specific stimuli. To obtain ERPs, we first segmented the continuous EEG signals into distinct epochs, locking at the onset of each word and extending 600 ms thereafter. This process resulted in a total of 4473 event-based epochs. Subsequently, we separately selected 50 trials out of 2924 trials for lexical words and 50 trials out of 1549 trials for functional words and calculated the average responses across all participants and all 64 channels over the epoch period (600ms), resulting in two event-based evoked data (Fig. 5D and E).

### Temporal signal-to-noise ratio (tSNR)

To quantify the strength of the signal at the voxel level, we computed tSNR both before and after preprocessing by dividing the mean signal intensity of a voxel across the time-series by its standard deviation. To assess the impact of extensive processing, we then compared the tSNR values before and after using Cohen’s d effect size:

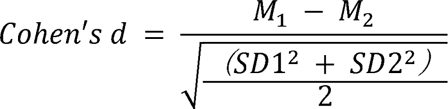

where M and SD are the mean and standard deviation of the tSNR in a voxel for the more minus the less preprocessed timeseries. The tSNR was computed for preprocessed functional runs in MNI152 space. We applied a gray matter mask with most white matter and ventricle voxels removed. The tSNR values showed a clear increase after denoising, suggesting a clearer signal compared to standard single echo acquisition (see Fig. 6).

**Fig. 6.**
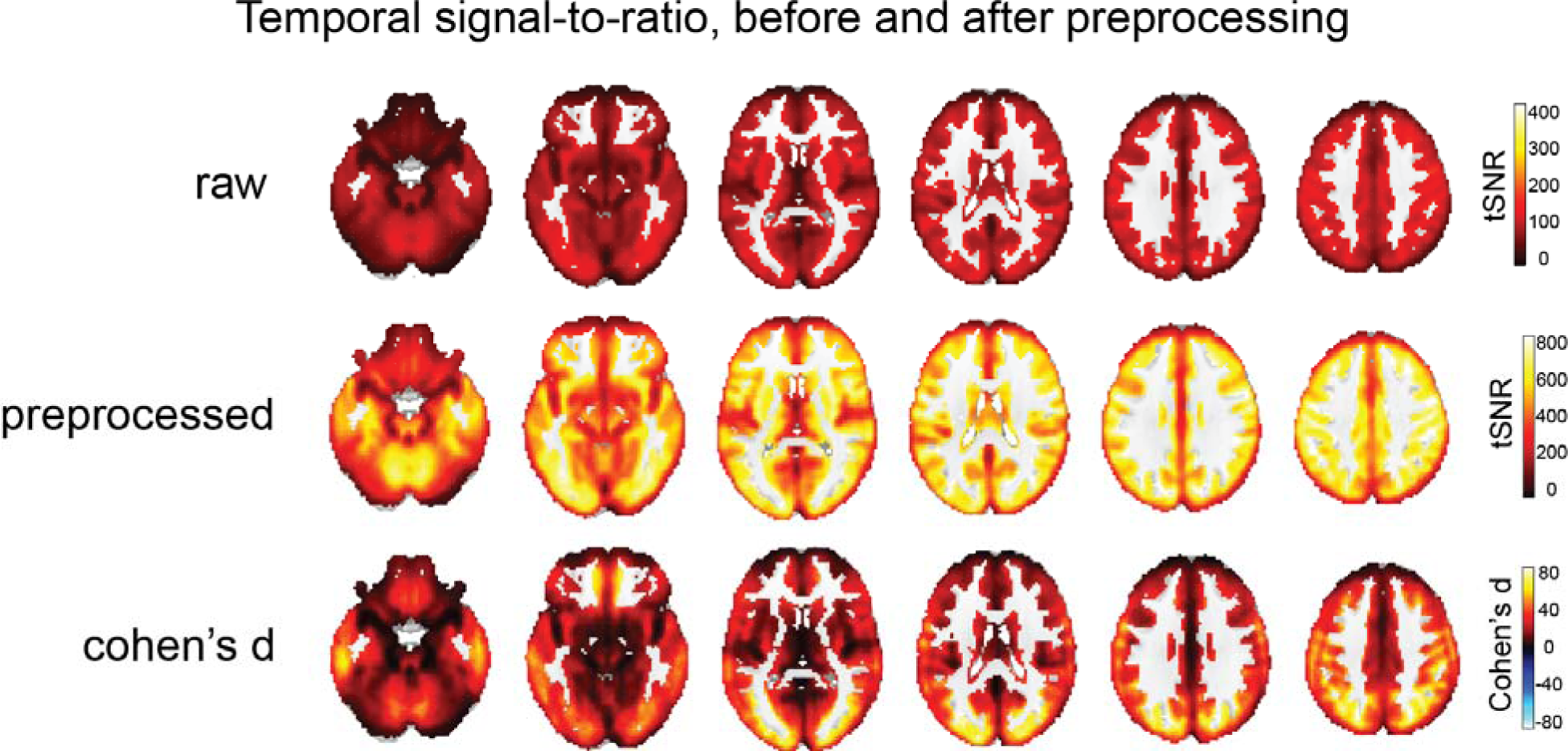
Voxel-wise temporal signal-to-noise ratio analysis before and after data preprocessing. Cohen’s d effect sizes showed an increase in tSNR after preprocessing

### Neural labeling

The general linear model (GLM) method was employed to derive the prosody- and word-relevant regions using our pitch and word annotations. We calculated the f0 and intensity for every 10 ms of the audio and marked the onset of each word in the audio (word rate). We then convolved the f0, intensity, and word rate annotations with a canonical hemodynamic response function and regressed them against the preprocessed fMRI timecourses using GLMs. At the group level, the contrast images for the f0, intensity and word rate regressors were examined by a one-sample t-test. Statistical significance was held at p < 0.05 FWE and the true positive was estimated by all-resolution inference^25^ (ARI). Figure 7 illustrates the GLM methods to localize the f0, intensity, and word rate regions. To illustrate the precise anatomical correspondence of our results with prior data, we overlaid fMRI term-based meta-analysis from Neurosynth (Retrieved October 2023) for the “pitch” areas (https://neurosynth.org/analyses/terms/pitch/; from 102 studies), the “acoustic” areas (https://neurosynth.org/analyses/terms/acoustic/; from 171 studies) and the “words’’ areas (https://neurosynth.org/analyses/terms/words/; from 948 studies). Our results are consistent with prior literature (see Fig. 8). Significant clusters from our GLM analysis and their statistics are shown in Supplementary Table 5.

**Fig. 7.**
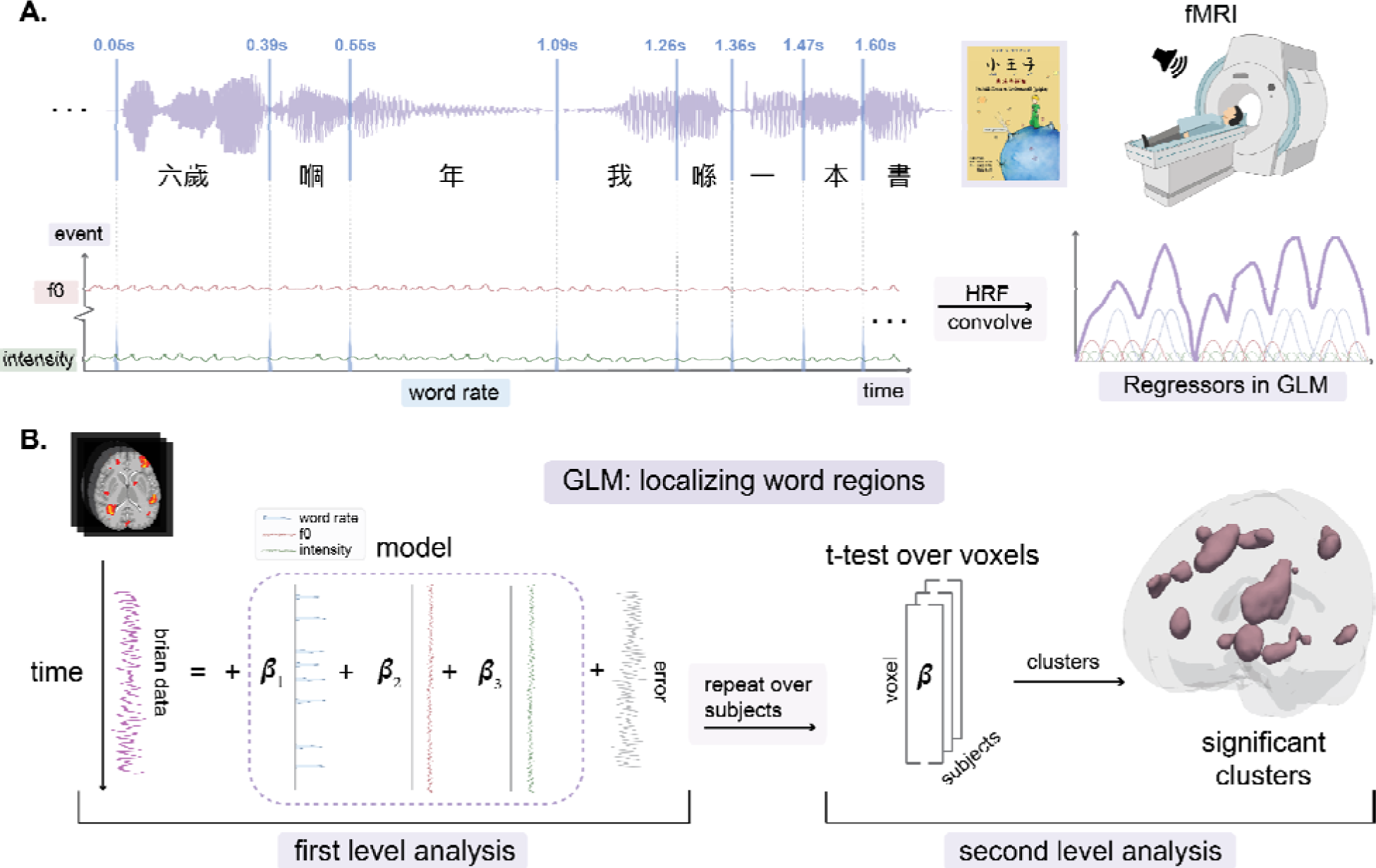
GLM analyses to localize the f0, intensity and word rate regressor. **(A)** Onset timepoints, f0 and intensity of each word in the audiobook were extracted and marked as 1 to be convolved with the canonical hemodynamic response function, separately. **(B)** At the first level, the time course of each voxel’s BOLD signals was modeled using the design matrix, including word rate, f0 and intensity as regressors. For the second-level analysis, a one-sample t-test was conducted on the distribution of the beta values for the included regressors across subjects at each voxel. The threshold of statistical significance was set at p < 0.05 FWE, k cluster > 50.

**Fig. 8.**
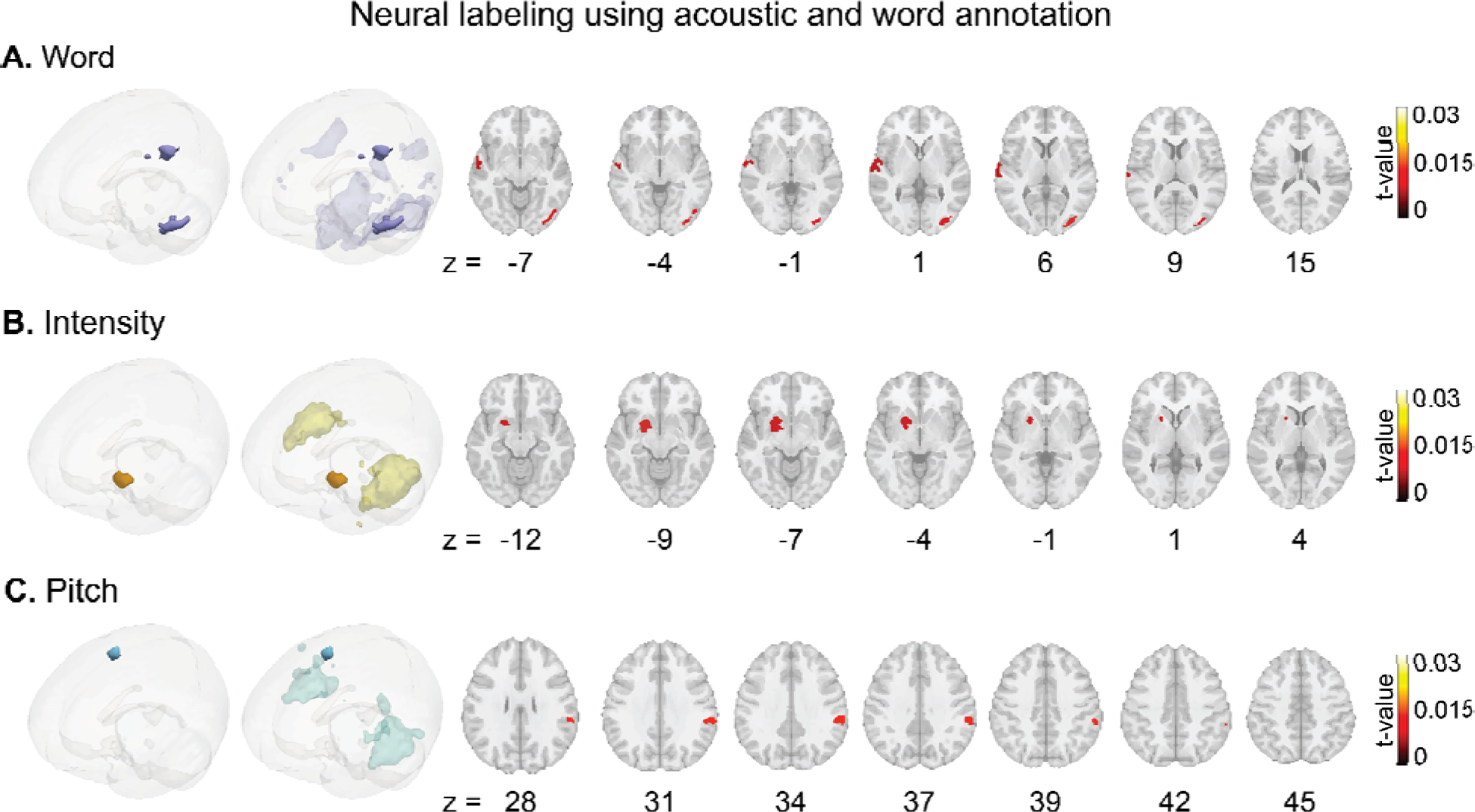
GLM results show the significant clusters for the **(A)** word, **(B)** intensity and **(C)** pitch regions using word rate, intensity and f0 annotations as regressors. In the second column of the 3D brain images, the light-colored areas represent related clusters obtained from meta-analyses of the word, acoustic, and pitch regions in Neurosynth. To determine statistical significance, a threshold of p < 0.05 FWE, k > 50 was applied.

## Usage Notes

The LPPHK fMRI-EEG dataset can advance our understanding of speech and language processing in the human brain, especially for older speakers, during naturalistic listening. However, there are several limitations and usage bottlenecks including annotations and analyses. We think discussing these limitations here can help others use the data more efficiently.

### Annotation bottleneck

Most linguistic annotations in our study were generated automatically utilizing the NLP tool, PyCantonese. However, a portion of the annotations required manual input from native Cantonese speakers. This manual aspect could lead to inconsistencies, potentially affecting subsequent annotations. Specifically, word segmentation tasks were conducted manually by two native speakers of Cantonese. Their manually segmented results were then used to ascertain word frequencies within the reference corpus, namely the Hong Kong Cantonese Corpus (Hkcancor). Notably, discrepancies between human and NLP- segmented results were observed, resulting in several words—707 out of 4473 total stimuli— exhibiting a zero frequency in the corpus.

### Analysis bottleneck

Although the GLM is widely employed in fMRI data analysis, there is a noticeable shift towards the adoption of encoding models in naturalistic paradigms. Nonetheless, there is still a need for accurate and systematic approaches to analyze complex, high-dimensional naturalistic fMRI data. Recent progress in machine learning-based methods and the integration of existing computational models has shown promise in improving compatibility with the intricate patterns of neural activity. Hence, we strongly encourage the development of advanced analysis approaches that take into account the various parameters of naturalistic stimuli. This will facilitate a more comprehensive exploration of the LPPHK fMRI-EEG dataset.

### Multi-functional analyses

LPPHK fMRI-EEG dataset is a valuable neuroscience resource since it combines neural data from two different modalities. We would like to propose several feasible research possibilities using this dataset. Firstly, owing to its remarkable spatial and temporal resolution, this dataset has the potential to elucidate the complex neural information processing and how it maps across various brain regions. As one part of the current proposed M/EEG-fMRI fusion analytical framework, this dataset can serve as a template for exploring further research possibilities. Secondly, as a dataset featuring an older age group, it offers a valuable resource for comparisons with other age groups, or even populations with specific clinical disorders. Finally, this database provides opportunities for a cross-linguistic analysis. For example, since Cantonese contains abundant inverted sentences and flexible word order, it provides valuable data which could be compared with previous databases such as LPPC-fMRI^23^.

## Code availability

The data for the LPPHK is publicly available at the OpenNeuro repository (https://openneuro.org/datasets/ds004718). The scripts can be accessed through GitHub (https://github.com/jixing-li/lpp_data).

## Acknowledgements

We want to thank Chou Man Jun, Chan Ka Man, Fung Wai Sze, Chun Wing Chan, and Ka Keung Lee for their help with data collection. We also want to thank PolyU Institute of Active Aging (now called the Research Centre for Gerontology and Family Studies) for helping us recruit participants. This work was supported by a Hong Kong Polytechnic University Start-up Fund (A0035856) given to MM.

## Contributions

MM, JB, JH, and LM designed the research, MM performed the research, MM, ZM, SW, CW and JL analyzed the data, MM, ZM, SW, CW and JL wrote the first draft, MM, JB, JH, LM and JL edited the paper.

---

**Supplementary Table 1.**
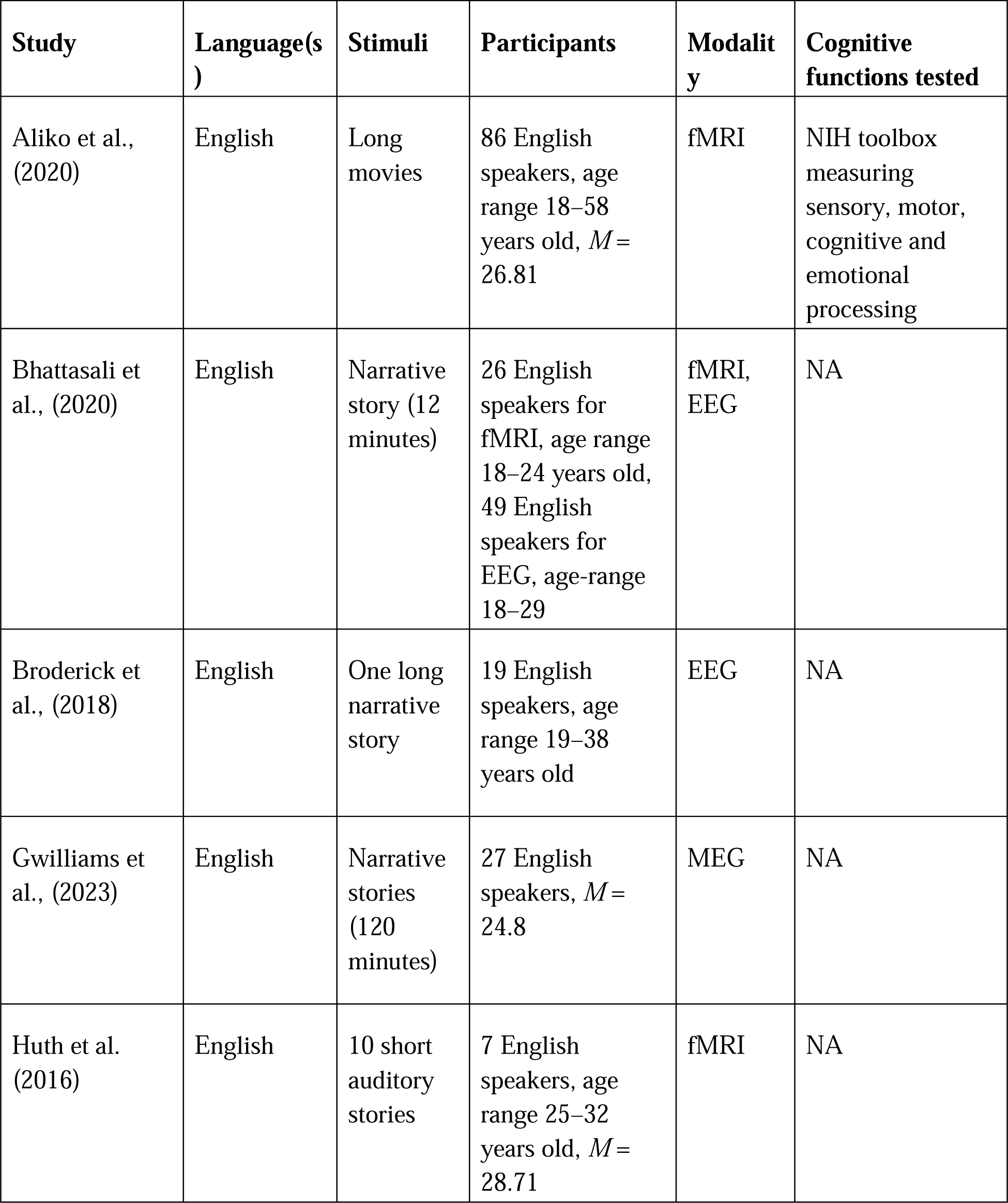
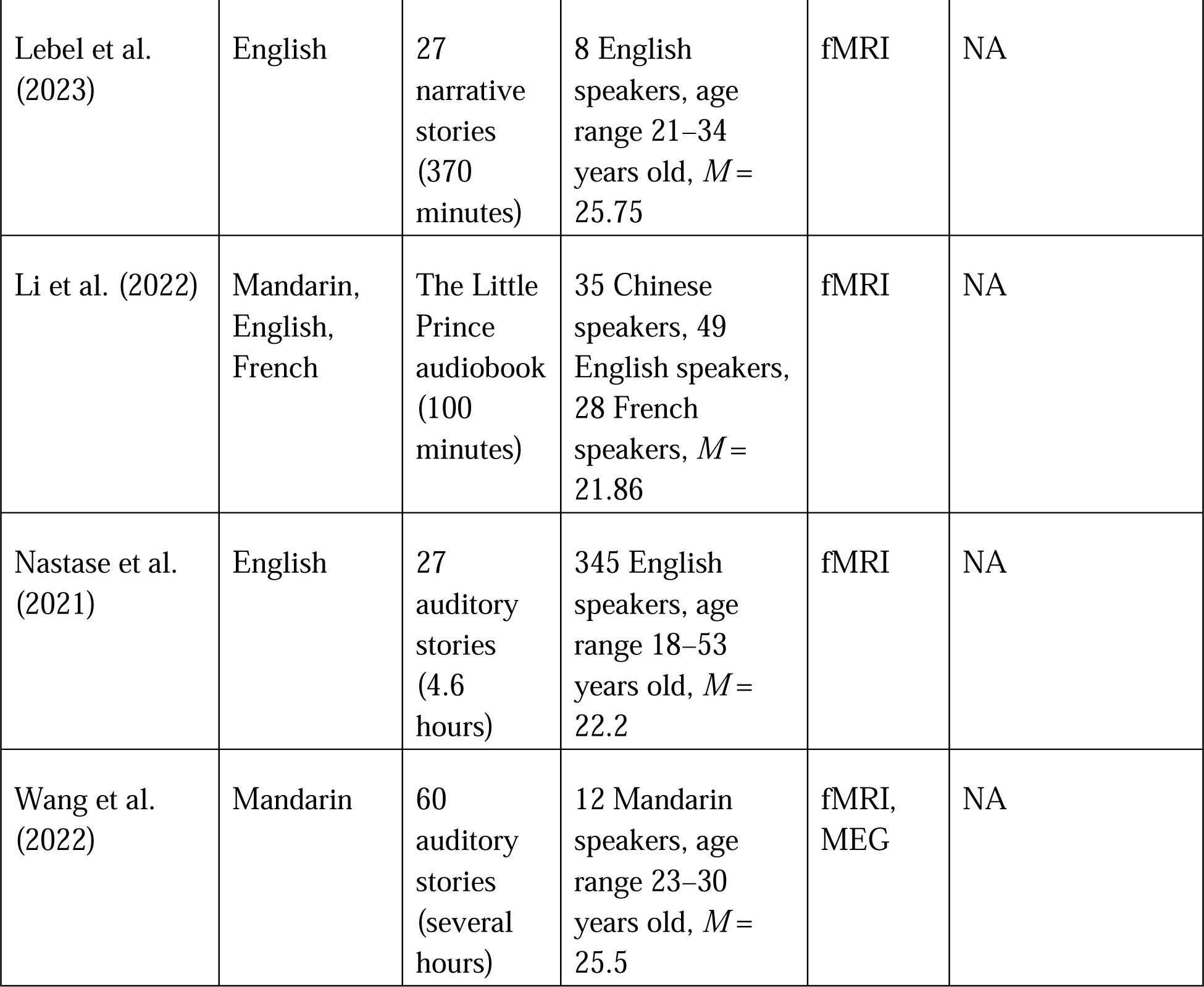
List of published naturalistic databases.

**Supplementary Table 2.** List of subjects in the data collection with basic demographic information.

| **Participants** | **Age** | **Sex** | **Education**<br>**(number**<br>**of years)** | **Languages**<br>**spoken** | **fMRI**<br>**data** | **EEG**<br>**data** | **Quiz**<br>**accuracy**<br>**of EEG\*** | **Quiz**<br>**accuracy**<br>**of fMRI\*** |
| --- | --- | --- | --- | --- | --- | --- | --- | --- |
| sub-HK001 | 65 | M | 9 | 1 | Y | Y | 16 | 13 |
| sub-HK002 | 67 | F | 9 | 1 | Y | Y | 10 | 11 |
| sub-HK003 | 71 | M | 9 | 2 | Y | Y | 13 | 13 |
| sub-HK004 | 72 | F | 12 | 2 | Y | Y | 12 | 10 |
| sub-HK005 | 76 | F | 6 | 1 | Y | Y | 11 | 7 |
| sub-HK006 | 66 | M | 9 | 2 | Y | Y | 12 | 13 |
| sub-HK007 | 66 | M | 9 | 2 | Y | Y | 16 | 16 |
| sub-HK008 | 77 | F | 9 | 2 | Y | Y | 14 | 8 |
| sub-HK009 | 77 | F | 6 | 2 | Y | Y | 10 | 10 |
| sub-HK010 | 77 | F | 6 | 1 | Y | Y | 9 | 4 |
| sub-HK011 | 67 | M | 9 | 2 | Y | Y | 12 | 12 |
| sub-HK012 | 67 | F | 9 | 3 | Y | Y | 14 | 12 |
| sub-HK013 | 66 | F | 9 | 3 | Y | Y | 16 | 10 |
| sub-HK014 | 67 | F | 6 | 2 | Y | Y | 13 | 10 |
| sub-HK015 | 65 | F | 6 | 1 | Y | Y | 13 | 13 |
| sub-HK016 | 71 | F | 12 | 4 | Y | Y | 15 | 13 |
| sub-HK017 | 69 | F | 9 | 1 | Y | Y | 18 | 11 |
| sub-HK018 | 66 | F | 9 | 2 | Y | Y | 16 | 11 |
| sub-HK019 | 69 | F | 6 | 2 | Y | Y | 11 | 15 |
| sub-HK020 | 69 | M | 9 | 3 | Y | Y | 11 | 10 |
| sub-HK021 | 65 | F | 12 | 3 | Y | Y | 14 | 11 |
| sub-HK022 | 65 | F | 9 | 3 | Y | Y | 14 | 9 |
| sub-HK023 | 68 | M | 9 | 4 | Y | Y | 11 | 10 |
| sub-HK024 | 72 | F | 9 | 2 | Y | Y | 10 | 9 |
| sub-HK025 | 66 | F | 9 | 4 | Y | Y | 11 | 15 |
| sub-HK026 | 67 | F | 9 | 3 | Y | Y | 18 | 15 |
| sub-HK027 | 68 | M | 16 | 3 | Y | Y | 15 | 15 |
| sub-HK028 | 67 | M | 12 | 1 | Y | Y | 11 | 12 |
| sub-HK029 | 69 | M | 9 | 3 | Y | Y | 12 | 14 |
| sub-HK030 | 65 | F | 9 | 4 | Y | Y | 14 | 14 |
| sub-HK031 | 67 | F | 12 | 2 | Y | N.A. | 15 | 13 |
| sub-HK032 | 66 | F | 9 | 3 | Y | Y | 14 | 11 |
| sub-HK033 | 72 | F | 6 | 3 | Y | Y | 9 | 11 |
| sub-HK034 | 65 | F | 6 | 4 | Y | Y | 15 | 13 |
| sub-HK035 | 66 | F | 9 | 2 | Y | Y | 16 | 13 |
| sub-HK036 | 71 | F | 6 | 2 | Y | Y | 12 | 11 |
| sub-HK037 | 69 | F | 6 | 2 | Y | Y | 16 | 12 |
| sub-HK038 | 71 | F | 9 | 2 | Y | Y | 10 | 14 |
| sub-HK039 | 67 | F | 12 | 3 | Y | Y | 16 | 16 |
| sub-HK040 | 65 | F | 9 | 2 | Y | Y | 14 | 10 |
| sub-HK041 | 70 | F | 16 | 3 | Y | Y | 10 | 11 |
| sub-HK042 | 69 | F | 9 | 1 | Y | Y | 16 | 12 |
| sub-HK043 | 70 | F | 6 | 2 | Y | Y | 10 | 12 |
| sub-HK044 | 75 | F | 9 | 2 | Y | Y | 9 | 13 |
| sub-HK045 | 71 | F | 6 | 1 | Y | Y | 12 | 13 |
| sub-HK046 | 69 | F | 16 | 3 | Y | Y | 17 | 14 |
| sub-HK047 | 74 | F | 6 | 2 | N.A. | Y | 15 | N.A. |
| sub-HK048 | 75 | F | 9 | 3 | N.A. | Y | 17 | N.A. |
| sub-HK049 | 66 | F | 9 | 2 | N.A. | Y | 8 | N.A. |
| sub-HK050 | 74 | M | 12 | 3 | N.A. | Y | 16 | N.A. |
| sub-HK051 | 69 | M | 12 | 1 | N.A. | Y | 14 | N.A. |
| sub-HK052 | 71 | F | 9 | 2 | N.A. | Y | 16 | N.A. |

\* Out of 20 questions

**Supplementary Table 3.** Cognitive tasks average data.

| Participants | Working memory | Picture Naming | Verbal Fluency | Flanker Task |  |  |
| --- | --- | --- | --- | --- | --- | --- |
|  |  |  |  | N_RT | IN_RT | C_RT |
| sub-HK001 | 7.5 | 1701.04 | 8.75 | 432.95 | 425.13 | 432.5 |
| sub-HK002 | 3 | 1528.60 | 7.00 | 524.47 | 535.39 | 537.23 |
| sub-HK003 | 2 | 1388.04 | 5.50 | 482.2 | 474.48 | 458 |
| sub-HK004 | 4 | 1422.10 | 11.50 | 568.37 | 569.4 | 552.52 |
| sub-HK005 | 3 | 1634.73 | 6.50 | 547.25 | 542.1 | 514.6 |
| sub-HK006 | 2.5 | 1488.99 | 10.25 | 412.78 | 431.02 | 387.48 |
| sub-HK007 | 3.5 | 1799.24 | 9.50 | 529.63 | 570.02 | 541 |
| sub-HK008 | 3 | 1783.09 | 7.00 | 521 | 576.89 | 508.62 |
| sub-HK009 | 3 | 1764.37 | 8.25 | 639.25 | 644.61 | 653 |
| sub-HK010 | 3 | 1566.61 | 9.50 | 577.46 | 581.72 | 520.5 |
| sub-HK011 | 1 | 1633.89 | 5.00 | 427.89 | 439.27 | 434.31 |
| sub-HK012 | 4 | 1731.08 | 10.00 | 509.28 | 475.82 | 481.08 |
| sub-HK013 | 4 | 1752.18 | 8.25 | 606.52 | 586.29 | 600.47 |
| sub-HK014 | 3 | 1689.51 | 10.50 | 485.6 | 524.76 | 488.62 |
| sub-HK015 | 3 | 1734.62 | 14.50 | 538.62 | 585.08 | 535.85 |
| sub-HK016 | 4.5 | 1620.30 | 11.00 | 532.81 | 581.36 | 505.71 |
| sub-HK017 | 4 | 1682.18 | 11.25 | 565.22 | 578.74 | 548.02 |
| sub-HK018 | 4.5 | 1485.14 | 9.25 | 437.61 | 491.25 | 435.47 |
| sub-HK019 | 4 | 1566.96 | 13.00 | 363.34 | 393.8 | 367.31 |
| sub-HK020 | 4 | 1421.30 | 12.00 | N.A. | N.A. | N.A. |
| sub-HK021 | 3.5 | 1345.61 | 12.25 | 406.97 | 430.16 | 404.22 |
| sub-HK022 | 4 | 1483.60 | 10.50 | 471.45 | 474.85 | 456.87 |
| sub-HK023 | 7 | 1458.89 | 11.25 | 435.5 | 451.86 | 408.97 |
| sub-HK024 | 4.5 | 1323.89 | 7.25 | 463.32 | 572.18 | 469.22 |
| sub-HK025 | 3.5 | 1279.41 | 12.00 | 420.94 | 470.27 | 418.1 |
| sub-HK026 | 4.5 | 1510.78 | 10.50 | 442.28 | 516.78 | 457.33 |
| sub-HK027 | 4 | 1886.45 | 13.75 | 395.3 | 447.5 | 427.87 |
| sub-HK028 | 4.5 | 1680.23 | 10.25 | 458.52 | 481.95 | 426.37 |
| sub-HK029 | 2 | 1252.21 | 13.00 | 387.19 | 377.95 | 371.88 |
| sub-HK030 | 3 | 1322.11 | 8.00 | 442.27 | 397.04 | 458.75 |
| sub-HK031 | 4 | 1610.27 | 13.50 | 447.95 | 437.94 | 438.68 |
| sub-HK032 | 4 | 1514.46 | 7.00 | 528.1 | 504.76 | 522.26 |
| sub-HK033 | 3.5 | 1135.66 | 10.50 | 472.61 | 468.43 | 488.9 |
| sub-HK034 | 5 | 1560.00 | 13.50 | 456.95 | 495.15 | 464.04 |
| sub-HK035 | 4 | 1765.84 | 13.25 | 506.35 | 505.4 | 490.27 |
| sub-HK036 | 3 | 1547.78 | 8.25 | 545.95 | 558.9 | 532.7 |
| sub-HK037 | 4 | 1753.05 | 6.50 | 551.47 | 540.37 | 545.08 |
| sub-HK038 | 3 | 1675.15 | 9.25 | 472 | 454.08 | 496.7 |
| sub-HK039 | 4.5 | 1700.43 | 11.00 | 484.25 | 444.5 | 456.68 |
| sub-HK040 | 6 | 1543.20 | 12.00 | 538.85 | 524.33 | 541.33 |
| sub-HK041 | 3 | 1567.36 | 10.25 | 398.06 | 431.52 | 448.89 |
| sub-HK042 | 3.5 | 1331.06 | 11.00 | 525.72 | 524.52 | 524.9 |
| sub-HK043 | 4.5 | 1328.60 | 11.25 | 497.5 | 468.05 | 485.25 |
| sub-HK044 | 4.5 | 1742.03 | 9.75 | 625.16 | 642.68 | 614.52 |
| sub-HK045 | 5 | 1516.42 | 7.75 | 512.96 | 528.53 | 536.8 |
| sub-HK046 | 6 | 1346.68 | 10.75 | 435.18 | 430.25 | 439.16 |
| sub-HK047 | 2.5 | 1778.82 | 6.25 | 668.54 | 648.37 | 644 |
| sub-HK048 | 3.5 | 1391.79 | 9.00 | 526.97 | 539.41 | 529.23 |
| sub-HK049 | 3.5 | 1461.17 | 7.75 | 544.54 | 559.28 | 540.91 |
| sub-HK050 | 3.5 | 1609.50 | 13.25 | 549.5 | 558.72 | 559.59 |
| sub-HK051 | 4 | 1488.49 | 12.75 | 631.58 | 652.32 | 637.2 |
| sub-HK052 | N.A. | N.A. | N.A. | N.A. | N.A. | N.A. |

$N_{RT}$ , refers to the reaction time from trials in the neutral condition. $IN_{RT}$ , refers to the reaction time from trials in the incongruent condition. $C_{RT}$ , refers to the reaction time from trials in the congruent condition.

**Supplementary Table 4.** Quality check for EEG data.

|  | **Bad Chan.** | **Epochs(4473)**<br>(Amplitude value,+/-150) |
| --- | --- | --- |
|  | **Pre-ICA** |  |
| sub-HK001 | None | 4464 |
| sub-HK002 | 1,2,3,4 | 4473 |
| sub-HK003 | 34,56,57 | 4461 |
| sub-HK004 | 1,2,3,4,5,30 | 4473 |
| sub-HK005 | None | 1333 (+/-150) , 3165(+/-300) |
| sub-HK006 | None | 4434 |
| sub-HK007 | None | 4460 |
| sub-HK008 | None | 4469 |
| sub-HK009 | 24,28 | 4456 |
| sub-HK010 | 4 | 4449 |
| sub-HK011 | None | 4473 |
| sub-HK012 | None | 4454 |
| sub-HK013 | None | 4522 |
| sub-HK014 | None | 4239 |
| sub-HK015 | 3 | 4473 |
| sub-HK016 | 3 | 3372 |
| sub-HK017 | None | 4324 |
| sub-HK018 | 37,46 | 4079 |
| sub-HK019 | None | 2182(+/-150) , 4060(+/-200) |
| sub-HK020 | 61 | 3409 |
| sub-HK021 | None | 4439 |
| sub-HK022 | None | 4452 |
| sub-HK023 | None | 4457 |
| sub-HK024 | None | 4320 |
| sub-HK025 | None | 4464 |
| sub-HK026 | None | 4470 |
| sub-HK027 | None | 4139 |
| sub-HK028 | None | 4463 |
| sub-HK029 | None | 0 (+/-150) , 3135 (+/-350) |
| sub-HK030 | None | 4437 |
| sub-HK032 | None | 4465 |
| sub-HK033 | None | 4465 |
| sub-HK034 | None | 4435 |
| sub-HK035 | None | 4420 |
| sub-HK036 | None | 4443 |
| sub-HK037 | None | 1251(+/-150), 3069(+/-200) |
| sub-HK038 | 3 | 4458 |
| sub-HK039 | 1,2,3,4 | 4473 |
| sub-HK040 | None | 4371 |
| sub-HK041 | 17 | 4190 |
| sub-HK042 | None | 4473 |
| sub-HK043 | None | 4223 |
| sub-HK044 | None | 4459 |
| sub-HK045 | None | 4465 |
| sub-HK046 | None | 4469 |
| sub-HK047 | None | 4468 |
| sub-HK048 | 29,30,31,38,40,41 | 4390 |
| sub-HK049 | None | 3221 |
| sub-HK050 | 34 | 4065 |
| sub-HK051 | 2,15 | 4470 |
| sub-HK052 | 19,27 | 4415 |

**Supplementary Table 5.** GLM results in MNI coordinate.

| Condition | Cluster | MNI coordinates | Cluster k-size (mm3) | *t* value | *p* value |
| --- | --- | --- | --- | --- | --- |
| Word rate | L Superior Temporal Gyrus | -57, -11, -10 | 127 | 6.41 | 0.006 |
|  | R Middle Occipital Gyrus | 48, -73, -10 | 120 | 6.44 | 0.001 |
| f0 | R Frontal Lobe | 53,-35,34 | 73 | 6.40 | 0.002 |
| Intensity | L Putamen | -27, 7, -10 | 128 | 7.01 | < 0.001 |

GLM results for the word rate, f0, and intensity regressors for fMRI data: MNI coordinates, cluster size and their peak level statistics, thresholded at $p < 0.05$ FWE and $k > 50$ .

